# Effects of Age and Noise Exposure History on Auditory Nerve Response Amplitudes: A Systematic Review, Study, and Meta-Analysis

**DOI:** 10.1101/2024.03.20.585882

**Authors:** James W. Dias, Carolyn M. McClaskey, April P. Alvey, Abigail Lawson, Lois J. Matthews, Judy R. Dubno, Kelly C. Harris

## Abstract

Auditory nerve (AN) function has been hypothesized to deteriorate with age and noise exposure. Here, we perform a systematic review of published studies and find that the evidence for age-related deficits in AN function is largely consistent across the literature, but there are inconsistent findings among studies of noise exposure history. Further, evidence from animal studies suggests that the greatest deficits in AN response amplitudes are found in noise-exposed aged mice, but a test of the interaction between effects of age and noise exposure on AN function has not been conducted in humans. We report a study of our own examining differences in the response amplitude of the compound action potential N1 (CAP N1) between younger and older adults with and without a self-reported history of noise exposure in a large sample of human participants (63 younger adults 18-30 years of age, 103 older adults 50-86 years of age). CAP N1 response amplitudes were smaller in older than younger adults. Noise exposure history did not appear to predict CAP N1 response amplitudes, nor did the effect of noise exposure history interact with age. We then incorporated our results into two meta-analyses of published studies of age and noise exposure history effects on AN response amplitudes in neurotypical human samples. The meta-analyses found that age effects across studies are robust (r=-0.407), but noise-exposure effects are weak (r=-0.152). We conclude that noise-exposure effects may be highly variable depending on sample characteristics, study design, and statistical approach, and researchers should be cautious when interpreting results. The underlying pathology of age-related and noise-induced changes in AN function are difficult to determine in living humans, creating a need for longitudinal studies of changes in AN function across the lifespan and histological examination of the AN from temporal bones collected post-mortem.

## Introduction

The auditory nerve (AN) is the only pathway connecting the inner ear to the central auditory system. It encodes acoustic information as differences in spike timing and transmission rates among neural fibers. Results of studies of AN pathology from animals and human temporal bones suggest that the AN is vulnerable to aging and noise exposure, more so than other cochlear structures (Fernandez et al., 2015; Kujawa & Liberman, 2015; Makary et al., 2011; Wu et al., 2019). The audiogram is the primary clinical tool for assessing hearing loss, but AN pathology can occur with or without measurable changes in pure-tone thresholds. For example, in chinchilla, only mild pure-tone threshold shifts were observed following carboplatin-induced 80% loss of inner hair cells and synaptic connections to AN fibers (Lobarinas et al., 2013; Takeno et al., 1994). The resulting loss of AN fibers would result in an under-sampling of the auditory signal (Lopez-Poveda & Barrios, 2013). While this under-sampling may not affect the detection of pure tones, it may disrupt the encoding of complex information, like speech. This can be especially problematic in noisy listening environments, where higher resolution of neural encoding is needed to preserve salient information within the acoustic signal. As such, loss or dysfunction of AN fibers is hypothesized to result in functional deficits that negatively impact speech-in-noise perception (Hickox & Liberman, 2014; Plack et al., 2014; Schaette & McAlpine, 2011).

Because pure-tone thresholds may not be a sensitive indicator of AN fiber loss, there is a need for clinically appropriate and non-invasive measures that are reliable indicators of AN function. In this pursuit, several non-invasive physiologic measures have been developed with the goal of assessing AN function in humans (for a review, see Harris & Bao, 2022). The sole non-invasive technique for directly assessing AN function in humans remains electroencephalography (EEG), specifically the measurement of Wave I of the auditory brainstem response (ABR WI) or the analogous N1 of the compound action potential (CAP N1). ABR WI and CAP N1 are potentials evoked by a brief auditory stimulus – measured from surface electrodes (ABR) or from the tympanic membrane (CAP) – and are thought to represent the action potentials at the distal AN (for a review, see Gibson, 2017). The capacity for these electrophysiologic measures to assess AN function is appealing because the methods for eliciting and measuring ABR WI and CAP N1 are already available in clinical settings. The accessibility of these techniques has contributed to a dramatic increase in the number of research studies employing EEG to assess AN function. This research has largely focused on two factors hypothesized to negatively impact AN function: 1) aging – age-related neurophysiologic changes in the AN, including synapse loss, AN fiber loss, and changes in AN myelin structure, can result in poorer AN function (e.g., Harris et al., 2021; Heeringa & Köppl, 2019; Kaewsiri et al., 2015; Konrad-Martin et al., 2012) – and 2) noise exposure – the noise-induced loss of cochlear synapses (cochlear synaptopathy) is thought to result in poor AN function in individuals with normal pure-tone thresholds, often referred to as ‘hidden hearing loss’ (e.g., Kim & Han, 2023; Plack et al., 2016).

AN function is largely comparable across mammalian species, with similar responses in rodents and humans (Burkard & Sims, 2001; Rumschlag et al., 2022). In rodent models, it is well-established that a loss of AN synapses and axons attributed to age or noise exposure can result in a significant decrease in ABR WI response amplitude (Kujawa & Liberman, 2015). Attempts to translate these contributing factors to specific aspects of AN function in humans, however, has generated mixed results (for reviews, see Bramhall et al., 2019; Le Prell, 2019; Ripley et al., 2022). Effects of age on AN response amplitudes are fairly consistent, with older adults exhibiting decreased ABR WI and CAP N1 amplitudes relative to younger adults (e.g., Burkard & Sims, 2001; Harris et al., 2021; Johannesen & Lopez-Poveda, 2021; but see Prendergast et al., 2019). Consistent with these electrophysiologic results from older humans, there is strong histological evidence from human temporal bones showing that aging contributes to AN pathology, specifically structural deficits in the AN, including a loss of cochlear synapses and AN fibers, as well as defects in the myelin structure of the remaining AN fibers (Makary et al., 2011; Nayagam et al., 2011; Wu et al., 2019; Xing et al., 2012). In contrast, associations between AN response amplitudes and a history of noise exposure (typically derived from self-report assessments) are mixed, with several studies reporting reduced ABR WI and CAP N1 response amplitude in individuals with a history of noise exposure (e.g., Bramhall et al., 2018a; Bramhall et al., 2017; Grose et al., 2017; Ridley et al., 2018; Skoe & Tufts, 2018), and others reporting a lack of significant associations (Fulbright et al., 2017; Grinn et al., 2017; Guest et al., 2019; Prendergast et al., 2019; Spankovich et al., 2017). Several factors have been identified that may contribute to discrepancies across these studies of noise exposure history, including the age and sex of participants and the methods used to assess noise exposure history and AN function (Le Prell, 2019). This is important because both increased age and noise exposure history are hypothesized to contribute to changes in ABR WI and CAP N1 response amplitudes. Most studies of noise-induced AN dysfunction have included participants in their 20s and 30s, few studies have included middle-aged and older adults, and among those, few test or control for age effects, and none test for interactions between the effects of age and noise exposure (Johannesen et al., 2019; Megha et al., 2021; Prendergast et al., 2019). A study of the effects of age and noise exposure history on AN response amplitudes within a single large sample can help disentangle how they each independently affect AN function and how they may interact to exacerbate AN dysfunction in older adults with a history of noise exposure.

### Current Investigation

To clarify the independent effects of age and noise exposure on electrophysiologic measures of AN function and determine how these two factors may interact, we tested the degree to which age and self-reported noise exposure history predict CAP N1 amplitudes in a large sample of younger (N=63) and older (N=103) adults. Results provide the first systematic examination of the interaction between age and noise exposure history on AN response amplitudes in human subjects. We then integrated the results of our study with those from prior studies in two meta-analyses; one to determine the mean effect of age and another the mean effect of noise exposure history on ABR WI and CAP N1 response amplitudes.

### Effects of Age and Noise Exposure History on CAP N1 response amplitudes Methods

#### Participants

CAPs were collected as part of several ongoing studies. Subsets of these data have been published elsewhere (Harris et al., 2021; McClaskey et al., 2018; Rumschlag et al., 2022), though associations of age and noise exposure history (described below) have not yet been reported in this larger sample of participants. CAPs were recorded in the right ears of 63 younger (18 to 30 years of age, mean (M)=23.9, standard deviation (SD)=3.2, 46 female) and 103 older (50 to 86 years of age, M=66.5, SD=7.2, 70 female) adults. Younger participants were graduate and undergraduate student volunteers from the Medical University of South Carolina and the College of Charleston in Charleston, SC. Older participants consisted of volunteers from the greater Charleston, SC area. Puretone audiometric thresholds for these participants are reported in **Figure 1**. The group of younger adults had audiometric thresholds ≤25 dB HL through 8 kHz. The group of older adults had audiometric thresholds that ranged from normal to mild-to-moderate sensorineural hearing loss, with more hearing loss in the higher test frequencies, more so for males than females. Self-reported noise exposure history was recorded in 58 younger and 85 older adults (Dubno et al., 2013). This study was approved by the Institutional Review Board of the Medical University of South Carolina and all participants provided written informed consent prior to participation.

**Figure 1.**
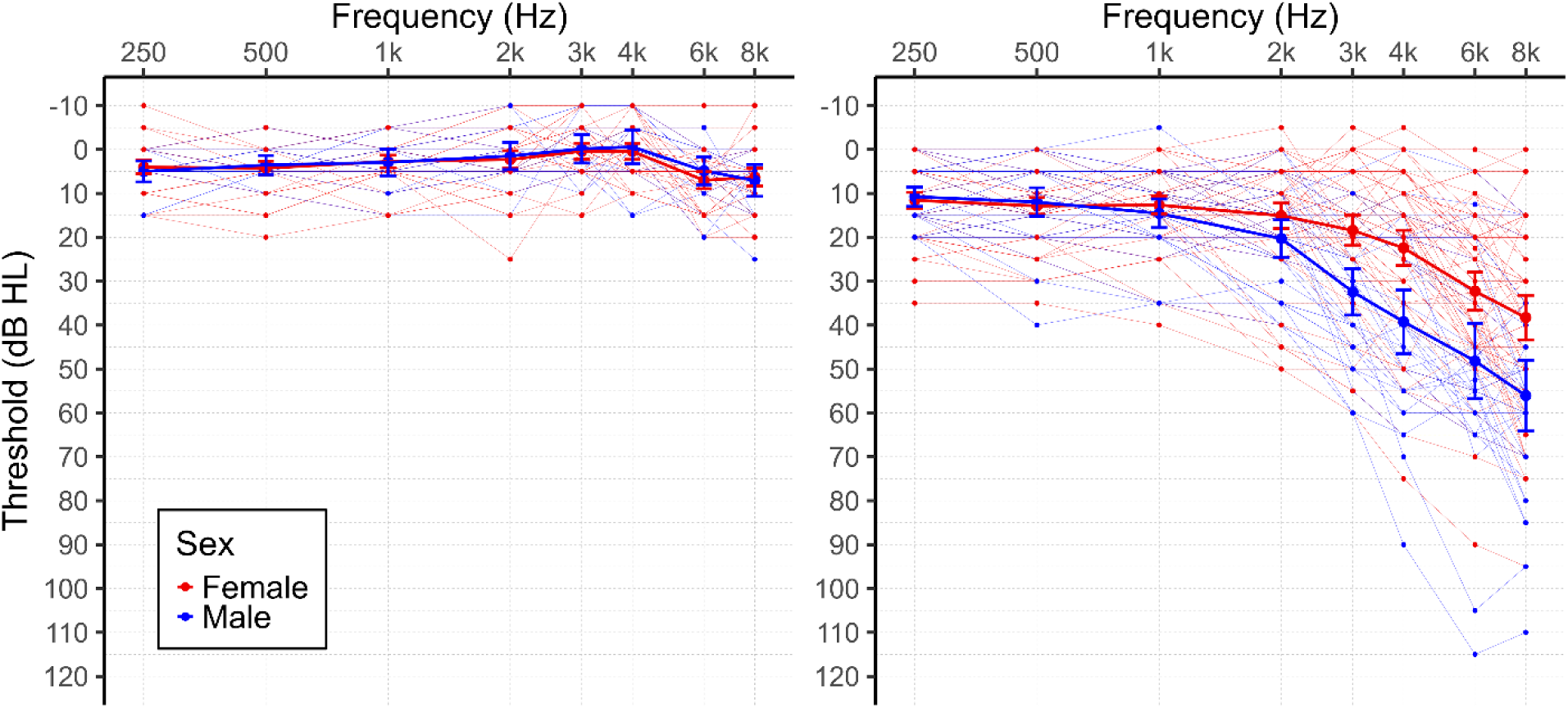
Pure-tone audiometric thresholds for the right (test) ear of individual younger (left) and older (right) participants. Red lines are female participants. Blue lines are male participants. Bold red and blue lines plot the mean pure-tone audiometric thresholds across female and male participants, respectively. Error bars indicate 95% confidence intervals.

#### Compound Action Potentials

CAPs were elicited by 100 microsecond alternating polarity rectangular pulses (clicks), presented at a rate of 11.1/s through an ER3C Insert Earphone (Etymotic Technologies) placed in the right ear. Stimuli and stimulus triggers were generated and controlled by RPvdsEx software from Tucker Davis Technologies (TDT). Stimuli were presented at 110 dB pSPL. CAPs were recorded from a tympanic membrane electrode (Sanibel) placed in the right ear, an inverting electrode placed on the contralateral (left) mastoid, and a low forehead grounding electrode. Auditory brainstem responses (ABRs) were simultaneously recorded (not reported here) for reference in identifying WI/N1 using a high forehead active electrode, an inverting mastoid electrode (on the right mastoid), and a low forehead grounding electrode. Responses were recorded in two blocks of 1100 trials (550 of each polarity) at a sampling rate of 20 kHz using a custom TDT headstage connected to the bipolar channels of a Neuroscan SynAmpsRT amplifier in AC mode with 2010x gain (Compumedics). Testing was done in an acoustically and electrically shielded booth. Participants reclined in a chair and were encouraged to rest quietly or sleep during recording.

Continuous AN activity was processed offline in MATLAB (MathWorks) using the EEGlab (Delorme & Makeig, 2004) and ERPLab (Lopez-Calderon & Luck, 2014) toolboxes. Continuous EEG signals were bandpass filtered between 0.150 and 3 kHz. The stimulus triggers sent by TDT RPvdsEx were shifted to account for the 1 ms delay introduced by the earphones and the 0.6 ms delay of the TDT digital-to-analog converter. The filtered data were epoched from -2 to 10 ms around stimulus onset and baseline corrected to a -2 ms to 0 ms prestimulus baseline (McClaskey et al., 2018).

Individual trials were rejected if peak deflections exceeded 45 μV using an automatic artifact detection algorithm and visual inspection. Epoched responses for the remaining trials were averaged. N1 peak selection was performed by two independent and experienced reviewers and assessed for repeatability across the two runs. Although only the 110 dB pSPL condition is reported here, CAP responses were collected from 70 to 110 dB pSPL to aid peak detection (e.g., Harris et al., 2022). The N1 peak-to-baseline amplitude and peak latency were measured in ERPlab using custom MATLAB functions.

#### Noise Exposure History Questionnaire

Noise exposure history was assessed using a seven-item noise history questionnaire on occupational and non-occupational noise exposure (**Supplemental Material**) (for details, see Dubno et al., 2013). Participants responded “yes” or “no” to whether they had been exposed to the following types of noise: occupational (including military), firearms, music, power tools, farm equipment, sudden loud noises, and other noises. Participants were identified as having a positive noise history if they reported exposure to at least one of the seven categories. Of the 58 younger and 85 older participants who completed the noise history questionnaire, 36 younger (25 female) and 51 older (26 female) reported a positive noise history. Evident from the distribution of responses were sex-related differences in noise exposure, especially for older participants. The proportion of older males who reported a positive noise exposure history (83%) was higher than the proportion of older females who reported a positive noise exposure history (47%).

#### Analytical Approach

We used linear regression models (LM) to test whether CAP N1 amplitudes were predicted by age group (younger or older) and noise exposure history (positive or negative). Participant sex was incorporated into our statistical analyses to evaluate whether sex-differences in AN response amplitudes and noise exposure history could account for variability in CAP N1 amplitudes explained by age and noise exposure history (e.g., Dewey et al., 2020; Fulbright et al., 2017; Grinn & Le Prell, 2022; Johannesen & Lopez-Poveda, 2021). Pure-tone thresholds were similarly incorporated into our statistical analyses to evaluate whether inter-subject variability in pure-tone thresholds across frequencies could account for variability in CAP N1 amplitudes explained by age and noise exposure history. We used two pure-tone threshold averages tested between different regression models. 1) Average pure-tone thresholds from 0.5 to 4 kHz were used to test for effects of hearing level within a frequency range often used to screen participants in studies of noise exposure history and AN function (e.g., Bramhall et al., 2017; Prendergast et al., 2017a; Stamper & Johnson, 2015). 2) Average pure-tone thresholds from 2 to 8 kHz were used to test for effects of hearing level within a frequency range where greater age-group and sex differences were observed (Figure 1), where energy from our broadband click stimulus is more concentrated, and where it has been theorized that noise-induced cochlear damage would be more prominent (e.g., Antonioli et al., 2016; Borchgrevink et al., 1996; Plack et al., 2016). Statistical analyses were performed in R (R Core Team, 2023).

### Results

Two LM models tested the degree to which CAP N1 response amplitudes are predicted by age and noise exposure history. Model 1 includes all participants and finds that CAP N1 response amplitudes are smaller (more positive) in older adults, compared to younger adults (B=0.065, SE_B_=0.030, β=0.339, SE_β_=0.158, t(164)=2.140, p=0.034). Adding participant sex to Model 1 did not improve model fit (χ^2^(1)=3.504, p=0.061) and participant sex was not a significant predictor of CAP N1 response amplitude after accounting for age-group differences. Additionally, Model 1 fit was not improved by adding pure-tone threshold averages from 0.5 to 4 kHz (χ^2^(1)=0.540, p=0.462) or from 2 to 8 kHz (χ^2^(1)=0.188, p=0.665) and neither average of pure-tone thresholds was a significant predictor of CAP N1 response amplitude after accounting for age-group differences. Model 2 includes the subset of participants with noise history data and found that noise exposure was not a predictor of CAP N1 response amplitudes (B=0.042, SE_B_=0.030, β=0.116, SE_β_=0.082, t(140)=1.402, p=0.163), though age was still a significant predictor (B=0.069, SE_B_=0.030, β=0.392, SE_β_=0.167, t(140)=2.341, p=0.021) (**Figure 2**). Adding participant sex to Model 2 did not improve model fit (χ^2^(1)=3.705, p=0.054), and participant sex was not a significant predictor of CAP N1 response amplitudes after accounting for age-group differences and noise exposure history. Additionally, Model 2 fit was not improved by adding pure-tone threshold averages from 0.5 to 4 kHz (χ^2^(1)<0.001, p=0.998) or from 2 to 8 kHz (χ^2^(1)=0.129, p=0.719) and neither average of pure-tone thresholds was a significant predictor of CAP N1 response amplitude after accounting for age-group differences and noise exposure history. Noise exposure did not interact with age, nor did the inclusion of the interaction term in Model 2 improve model fit, χ^2^(1)=0.110, p=0.740.

**Figure 2.**
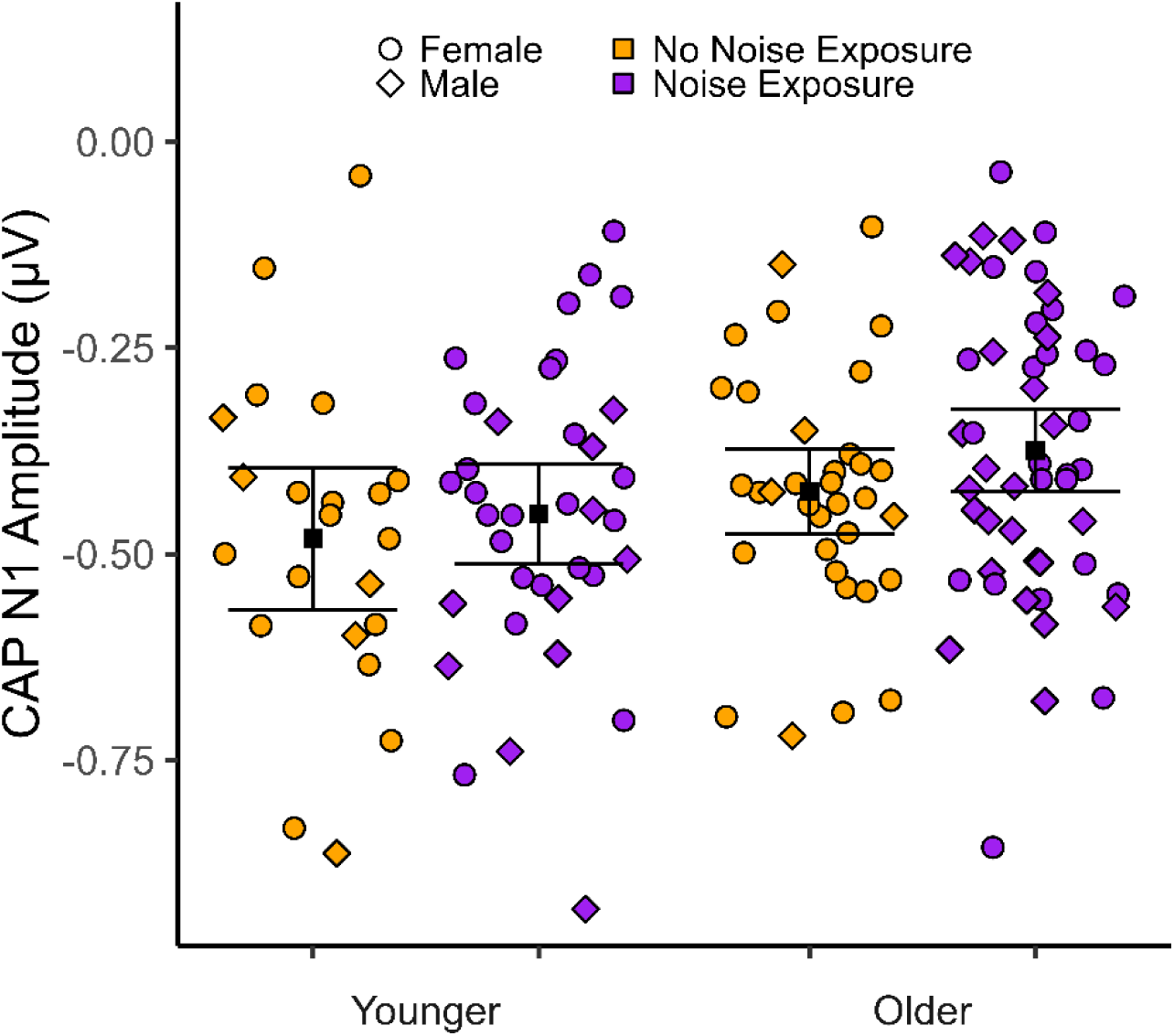
Dot plot of the CAP N1 amplitudes of younger (left) and older (right) participants. Circles indicate female participants. Diamonds indicate male participants. Participants with a history of noise exposure are marked in purple and participants with no history of noise exposure are marked in orange. Black boxes mark the group mean. Error bars indicate the 95% confidence intervals for the mean of each group.

### Summary

The results suggest that CAP N1 response amplitudes were smaller in older than younger adults. In contrast, CAP N1 response amplitudes were not predicted by self-reported history of noise exposure after accounting for participant age. Importantly, the relationship between noise exposure history and CAP N1 response amplitudes did not differ between younger and older participants. There has been some evidence suggesting that not all types of noise exposure affect the AN equally. Impulse/impact noise, such as from firearms, has been found to be negatively associated with AN response amplitudes (e.g., Grinn & Le Prell, 2022; Pinsonnault-Skvarenina et al., 2022b). When our data were reanalyzed with noise exposure exclusively restricted to a history of exposure to firearms, the pattern of relationships reported above did not change. We note, however, that our participants who reported a positive history of firearms exposure also largely reported using hearing protection.

Audibility is another factor to consider in studies of AN function. We controlled for effects of audiometric thresholds statistically by incorporating average pure-tone threshold from 0.5 to 4 kHz and from 2 to 8 kHz as covariates in our analyses. We found that pure-tone thresholds were not significant predictors of CAP N1 response amplitudes after accounting for age group. Alternatively, some studies control for audibility by restricting their sample to participants with normal hearing below 4 kHz, which may control for other pathologies that may affect AN function. Many studies that attribute differences in AN response amplitudes between noise-exposed and non-noise-exposed groups to noise-induced cochlear synaptopathy and “hidden hearing loss” have restricted their samples to those with normal hearing in this way (e.g., Bramhall et al., 2017; Prendergast et al., 2017a; Stamper & Johnson, 2015). To replicate these methods and more thoroughly control for cochlear pathologies that may underlie hearing loss and affect AN function, we reanalyzed our data in only those younger and older participants with pure-tone thresholds ≤25 dB HL at 0.5, 1, 2, 3, and 4 kHz. The pattern of relationships reported above did not change when our analyses included CAP N1 response amplitudes from only those participants, suggesting that statistical controls for audiometric thresholds were sufficient to account for the effects of audibility and that our results are not driven by additional pathologies that are comorbid with elevated audiometric thresholds.

Failing to find support for a hypothesized relationship between noise exposure history and AN function in our large cohort of participants is consistent with many past studies (Fulbright et al., 2017; Grinn et al., 2017; Guest et al., 2019; Johannesen et al., 2019; Prendergast et al., 2019; Spankovich et al., 2017). Our results suggest that studies finding positive noise history to negatively impact AN function in humans may be explained by uncontrolled factors, such as methodology and sample demographics, or they may capture a relationship that is small across the broader population.

### AN Function as Predicted by Age and Noise-Exposure History: Meta-Analyses

To further examine the relationship between AN function and age and between AN function and noise exposure, we performed two meta-analyses to measure these relationships across the broader literature.

### Methods

#### Studies

We first conducted a PubMed advanced search of key terms to find studies of ABR WI and CAP N1 in neurotypical adult human samples. For ABR studies, the advanced search included the terms “auditory brainstem responses” and “wave I”, excluded the terms “infants”, “children”, “schizophrenia”, “ADHD”, and “autism”, and were filtered to include only human studies. For CAP studies, the advanced search included the terms “compound action potential” and “auditory nerve”, excluded the term “eCAP”, and were filtered to include only human studies. The large number of studies returned by these search queries (1,278 ABR, 223 CAP) were checked by three experts (authors Dias, Harris, and McClaskey) to confirm they matched our search criteria and further filtered to include only those studies that measured ABR WI or CAP N1 response amplitude and explicitly tested and reported the relationship between ABR WI or CAP N1 response amplitude and age or noise exposure. As a consequence, some studies that motivated our study were excluded from one or both meta-analyses for reporting the relationships of other metrics of AN function instead of ABR WI/CAP N1 response amplitude (e.g., Gómez-Álvarez et al., 2023; Johannesen et al., 2019) or for not providing descriptive or inferential statistics needed to calculate the effect size of the relationship between ABR WI/CAP N1 response amplitude and age or noise exposure history, typically for non-significant relationships (e.g., Maele et al., 2021; Megha et al., 2021; Prendergast et al., 2019; Ripley et al., 2022). The latter is important to consider because the mean effect size computed across studies may be biased by significant effects reported within published studies, providing an inflated mean effect size that is larger than the true population mean (Cumming, 2012; Rosenthal, 1991). Of the studies that met our criteria, a citation search was performed in PubMed and Google Scholar for the most cited studies (e.g., Prendergast et al., 2017a; Stamper & Johnson, 2015) to find any other studies that may have met our inclusion criteria but were otherwise missed by the PubMed search. These procedures were completed on March 5, 2024, and resulted in 20 studies (14 different study sites) of age and ABR-WI/CAP-N1 amplitude and 27 studies (20 different study sites) of noise exposure and ABR-WI/CAP-N1 amplitude. We then incorporated our own CAP data reported in the previous section (listed as “Dias et al., 2024”) into this pool of studies for our meta-analyses. The final number of studies included in our meta-analysis of age and ABR-WI/CAP-N1 amplitude was 21 (14 different study sites) and the final number of studies included in our meta-analysis of noise exposure and ABR-WI/CAP-N1 amplitude was 28 (21 different study sites). When collecting studies for our systematic review and meta-analysis, we used conventional best practices as recommended by the Preferred Reporting Items for Systematic Reviews and Meta-Analysis (PRISMA) statement (Page et al., 2021a; Page et al., 2021b) and others (Cumming, 2012; Rosenthal, 1991).

#### Analytical Approach

Separate meta-analyses were performed to measure the average (mean) relationship between ABR-WI/CAP-N1 amplitude and age, and the relationship between ABR-WI/CAP-N1 amplitude and noise exposure history. For each study included in these meta-analyses, we computed the effect size (r_Effect Size_) for each relationship between response amplitude and age or between response amplitude and noise exposure history from the descriptive and inferential statistics reported in each study. Studies often reported these relationships for multiple different conditions, which can vary by participant characteristics (sex, age, and noise exposure), ear tested, electrode configuration (surface and canal electrode configurations for ABRs and tympanic-membrane electrode configurations for CAPs), and stimulus characteristics (stimulus type, duration, intensity, and presentation rate). As such, we often extracted multiple values from within individual studies. All r-values were transformed into Z-scores using Fisher’s transformation. For studies of ABR-WI/CAP-N1 response amplitude and age, if response amplitude decreased as age increased, then the Z-score was coded as negative and otherwise coded as positive. For studies of ABR-WI/CAP-N1 response amplitude and noise exposure history, if response amplitude decreased with more noise exposure history, then the Z-score of the relationships was coded as negative and otherwise coded as positive. These Z-scores were used as the outcome variables of our meta-analyses (e.g., Dias et al., 2022; Hansen et al., 2022; Rosenthal, 1991; Rosnow et al., 2000).

To account for the nestedness of the multiple relationships reported in many individual studies and to account for the nestedness of many studies that were conducted at the same institution (typically by the same research group), linear mixed effects regression (LMER) was used to conduct our meta-analyses. Accounting for the nestedness of studies within study sites is important because investigators often report results from overlapping samples to test different hypotheses in different studies. Different LMER models were used to perform separate meta-analyses for studies of ABR-WI/CAP-N1 response amplitude and age and for studies of ABR-WI/CAP-N1 response amplitude and noise exposure history. These LMER models included the Z transforms of relationship effect sizes as the outcome variable and study and study site as random factors (grouping variables). Each study was weighted by their reported sample size. Weighting studies by sample size is important when calculating the mean effect across a pool of studies (for a review, see Cumming, 2012), but we recognize that the sample sizes of our CAP study is disproportionate to most other studies in our meta-analyses. To ensure that the inclusion of our CAP study did not significantly bias the mean effect calculated across studies, we performed a Z-test of the difference in effects between meta-analyses with our CAP study and without:

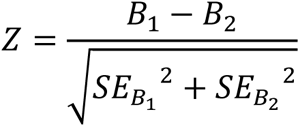

where *B_1_* is the coefficient (mean Z-transformed effect across studies) for the meta-analysis with all studies and *B_2_* is the coefficient for the meta-analysis with all studies except our current CAP study. *SE_B_1__* and *SE_B_2__* are the standard errors for the respective coefficients (Clogg et al., 1995; Cohen et al., 2003; Dias et al., 2021; Dias et al., 2022; McClaskey et al., 2018; Paternoster et al., 1998).

We also considered whether electrode configuration (surface, canal, or TM), stimulus characteristics (stimulus, stimulus duration, stimulus intensity, and presentation rate), or sample sex (performed only in the subset of nine studies that reported effects of noise exposure history separately for male and female participants – too few studies report effects of age separately for male and female participants) modulated the relationships of interest in our meta-analyses, but none were found to be significant predictors (p>0.2) and their inclusion in the LMER models did not improve model fit (p>0.2).

Meta-analyses were performed using the *meta* (Balduzzi et al., 2019) and *metafor* (Viechtbauer & Cheung, 2010) packages for *R* (Harrer et al., 2021; R Core Team, 2023).

### Results

The meta-analysis of the relationship between ABR-WI/CAP-N1 response amplitude and age found that the mean effect across studies is significant (B=-0.432, SE=0.127, t=-3.393, p=0.002). The forest plot in **Figure 3** illustrates the Fisher Z-transforms of the relationships reported in each study and the mean Z-transformed relationship between ABR-WI/CAP-N1 amplitude and age across studies, which when reverse transformed equates to a mean effect of r=-0.407, 95% CI [-0.600, -0.169].

**Figure 3.**
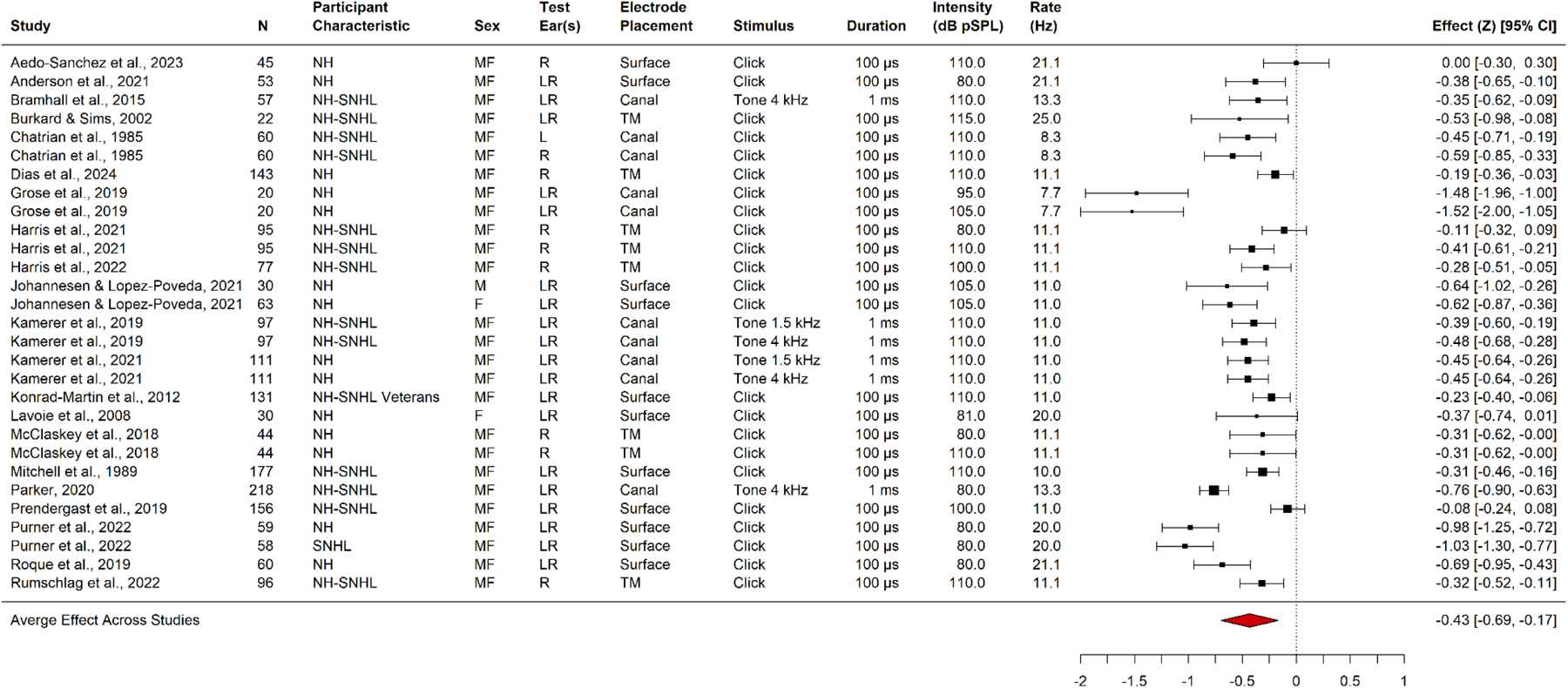
Forest plot of the studies included in our meta-analysis of the relationship between age and AN response amplitude. The right columns report the magnitude and 95% confidence interval of the Z-transformed effect size for each experimental group within each study. The size of the box for the effect of each study represents the study sample size. The mean effect and 95% confidence interval across studies is represented by the red diamond. For “Participant Characteristics”, “NH” indicates the sample was comprised of adults with normal hearing, “SNHL” indicates the sample was comprised of adults with sensorineural hearing loss, and “NH-SNHL” indicates the sample was comprised of adults with normal hearing or sensorineural hearing loss. For “Sex”, “M” indicates the sample was comprised of only males, “F” indicates the sample was comprised of only females, and “MF” indicates the sample was comprised of both males and females. For “Test Ear(s)”, “L” indicates testing was performed only in the left ear, “R” indicates testing was performed only in the right ear, and “LR” indicates testing was performed in either or both ears. “Electrode Placement”, indicates where the active electrode was placed for recording of the AN response, either outside of the ear (“Surface”), in the ear canal (“Canal”), or on the tympanic membrane (“TM”).

Response amplitudes of ABR WI and CAP N1 are smaller in older than in younger adults. The mean effect did not significantly change when removing our current CAP study from the meta-analysis, Z=0.092, p=0.464.

The meta-analysis of the relationship between ABR-WI/CAP-N1 response amplitude and noise exposure history found that the mean relationship across studies is significant (B=-0.153, SE=0.043, t=-3.562, p<0.001). The forest plot in **Figure 4** illustrates the Fisher Z-transforms of the relationships reported in each study and the mean Z-transformed relationship between ABR-WI/CAP-N1 response amplitude and noise exposure history across studies, which when reverse transformed equates to a mean effect of r=-0.152, 95% CI [-0.233, -0.068]. ABR WI and CAP N1 response amplitudes are smaller in individuals with a positive noise exposure history. The mean effect did not change when removing our current CAP study from the meta-analysis, Z=0.016, p=0.494.

**Figure 4.**
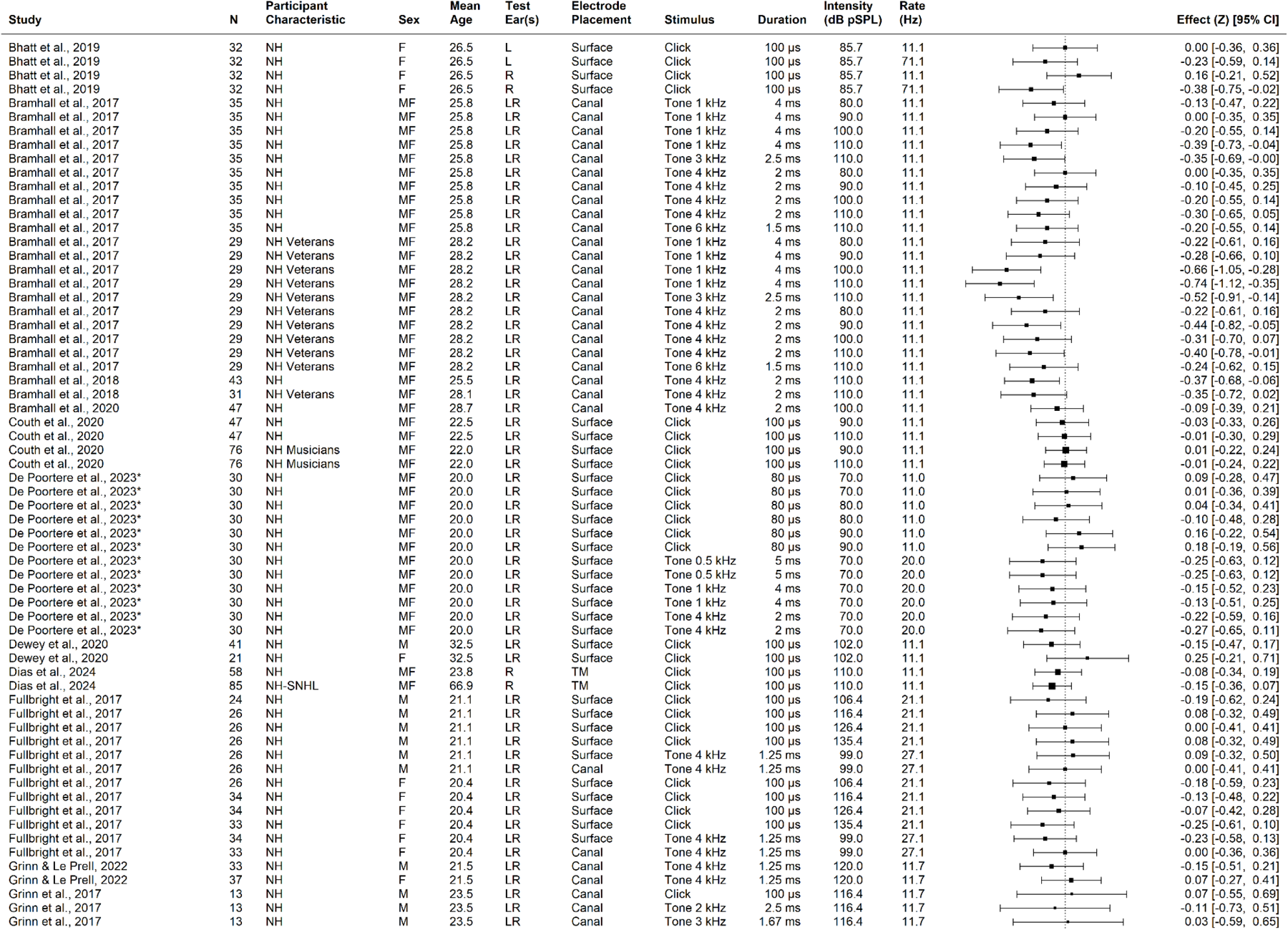
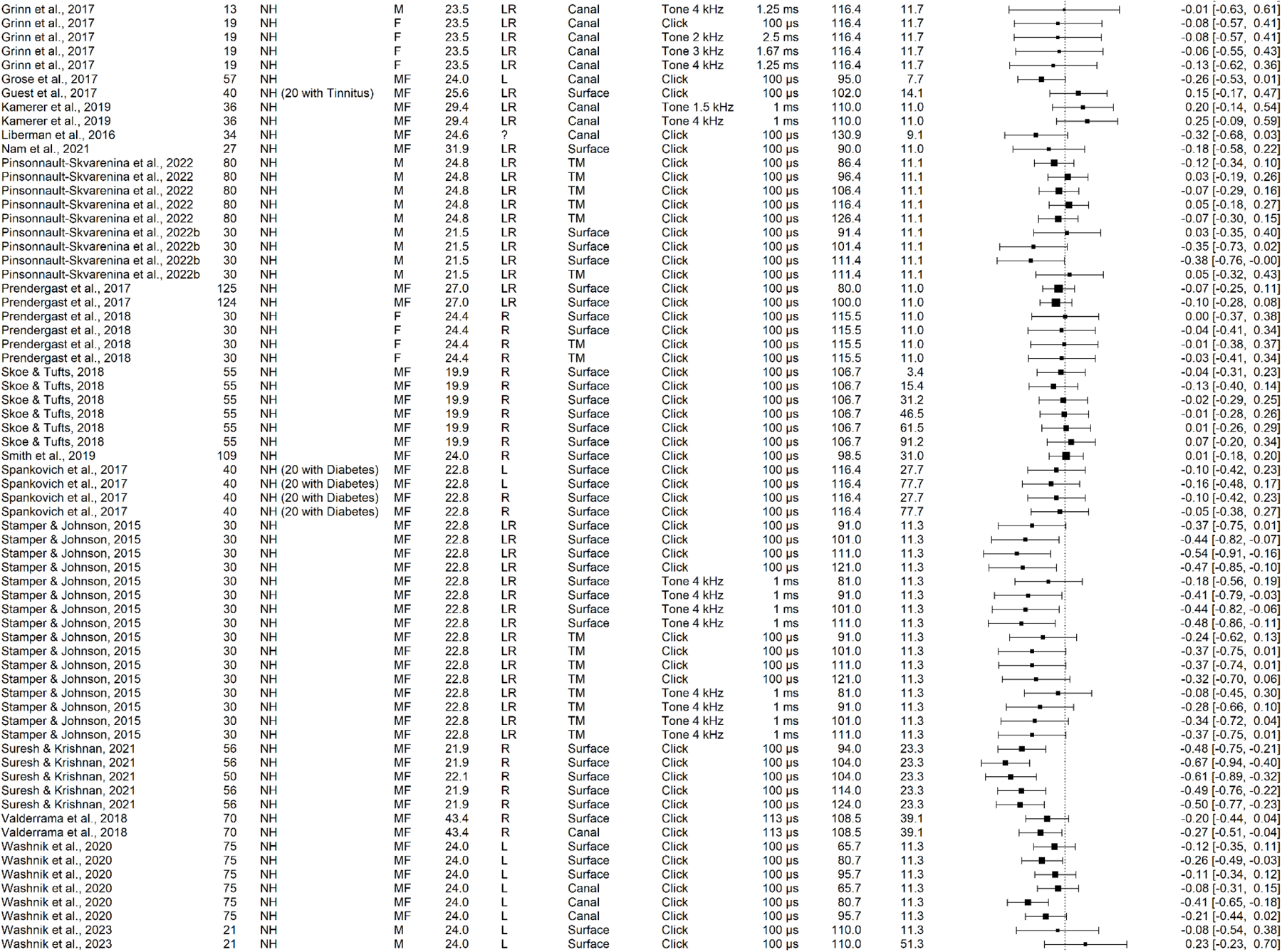
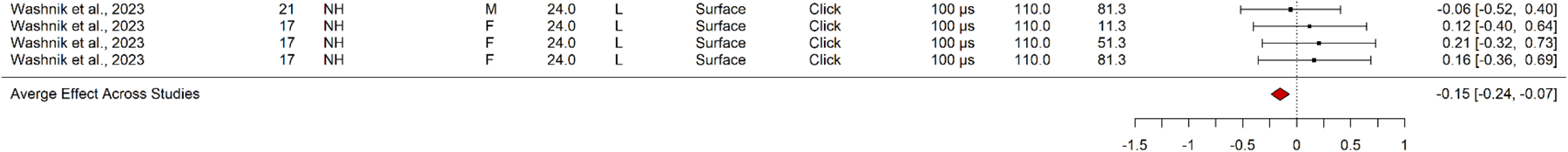
Forest plot of the studies included in our meta-analysis of the relationship between noise exposure history and AN response amplitude. The right columns report the magnitude and 95% confidence interval of the Z-transformed effect size for each experimental group within each study. The size of the box for the effect of each study represents the study sample size. The mean effect and 95% confidence interval across studies is represented by the red diamond. For “Participant Characteristics”, “NH” indicates the sample was comprised of adults with normal hearing, “SNHL” indicates the sample was comprised of adults with sensorineural hearing loss, and “NH-SNHL” indicates the sample was comprised of adults with normal hearing or sensorineural hearing loss. For “Sex”, “M” indicates the sample was comprised of only males, “F” indicates the sample was comprised of only females, and “MF” indicates the sample was comprised of both males and females. For “Test Ear(s)”, “L” indicates testing was performed only in the left ear, “R” indicates testing was performed only in the right ear, and “LR” indicates testing was performed in either or both ears. “Electrode Placement”, indicates where the active electrode was placed for recording of the AN response, either outside of the ear (“Surface”), in the ear canal (“Canal”), or on the tympanic membrane (“TM”). *De Poortere et al., 2023 used two different self-report measures of noise exposure history and so reported two relationships between noise exposure history and ABR WI amplitude for each stimulus-intensity combination they tested.

### Summary

The meta-analyses found that both age and noise exposure history are predictors of AN response amplitudes across studies. ABR-WI/CAP-N1 response amplitudes decline with age and with noise exposure history. Importantly, the mean effects of age and noise exposure history calculated across studies were not biased by the inclusion of our larger CAP study. The small mean effect size of the relationship between noise exposure history and AN response amplitude is not surprising, given the number of individual studies that failed to find support for a relationship between noise exposure history and AN response amplitudes, including our own. It should be considered that the meta-analyses included only published studies, subject to publication bias, and consideration should be given to ‘file-drawer’ studies that never made it to publication (Rosenthal, 1979; Rosenthal, 1991).

## Discussion

The goal of the systematic review, study, and meta-analyses reported here was to determine the extent to which age and noise exposure history impacted AN function. We focused on the most often reported aspect of AN function: suprathreshold ABR WI and CAP N1 response amplitudes. We first examined the extent to which age and noise exposure history predict CAP N1 response amplitude in a large cohort of younger and older adults. Our results suggest that CAP N1 response amplitudes decline with age, but noise exposure history was unrelated. Although prior animal studies have suggested an interaction between noise exposure and age, with a greater decrease in AN function in aged animals with prior noise exposure than those without noise exposure (Fernandez et al., 2015), the interaction between age and noise exposure history was not a significant factor in our CAP study. The meta-analyses found that ABR WI/CAP N1 response amplitudes are smaller in older than younger adults and in adults with a positive rather than negative noise exposure history. While the relationship between age and AN function is fairly conclusive, with a robust effect across studies (r=-0.407, **Figure 3**), the association between noise exposure history and AN response amplitudes is weak (r=-0.152, **Figure 4**), requiring further discussion of how to interpret effects of noise exposure on AN function.

### Effects of Age on AN Response Amplitudes

Significant effects of age on AN response amplitudes were observed in our CAP study and in our meta-analysis, adding to a growing literature suggesting that AN function declines with age. Findings from animal models suggest that aging accelerates AN damage associated with environmental exposures (Fernandez et al., 2015). Despite these findings in animals, few studies have tested both the effects of age and noise exposure on AN function in humans (Johannesen et al., 2019; Megha et al., 2021; Prendergast et al., 2019). Unfortunately, these studies did not include participants over the ages of 60 (Prendergast et al., 2019), 65 (Megha et al., 2021), and 68 (Johannesen et al., 2019). The results are also conflicting between studies, with one study finding an effect of both age and noise exposure history (Megha et al., 2021), another finding no effect of either age or noise exposure (Prendergast et al., 2019), and the last study finding an effect of age, but not noise exposure (similar to our own CAP study) (Johannesen et al., 2019). Critically, none of these studies examined how an interaction between age and noise exposure history may relate to AN function. In our CAP study, we did not find a significant interaction between age and noise exposure history. Even though we found that CAP N1 response amplitudes were smaller in older than younger adults, a positive noise exposure history did not exacerbate age-related deficits in AN function.

### Effects of Noise Exposure History on AN Response Amplitudes

One goal of the present study was to resolve the often-conflicting reports regarding the effects of noise exposure history on the AN (for reviews, see Bramhall et al., 2019; Le Prell, 2019; Ripley et al., 2022). Evident across the literature are many different methods for eliciting and recording responses from the AN (**Figure 4**), which can contribute to the variability in results seen among studies. Nevertheless, the meta-analysis did not find evidence suggesting that the relationship between noise exposure and AN response amplitudes reported across the literature varied by the methods used to elicit (stimulus, duration, intensity, and presentation rate) or record (surface and canal electrodes for ABRs and TM electrodes for CAPs) AN responses.

It is important to consider not just the methods for eliciting and recording AN responses, but also to consider the composition of the study sample. Sex differences in self-reported noise exposure history and AN response amplitudes have been reported in other studies, with males more likely to report a positive noise exposure history and exhibit smaller AN response amplitudes than females (e.g., Dewey et al., 2020; Fulbright et al., 2017; Grinn & Le Prell, 2022; Johannesen et al., 2019). Previous studies of noise exposure history and AN function that have investigated and controlled for sex differences in the relationship between noise exposure and AN response amplitudes have failed to identify a significant relationship between noise exposure history and AN function in either males or females (Dewey et al., 2020; Fulbright et al., 2017; Grinn & Le Prell, 2022; Grinn et al., 2017; Washnik et al., 2023). We also did not find evidence for a difference in the relationship between noise exposure history and AN response amplitudes between male and female participants in our meta-analysis. It is possible that the negative relationship between noise exposure history and AN function reported in some studies that do not control for participant sex can be explained by a sample comprised of more males with higher rates of positive noise exposure histories and smaller ABR WI/CAP N1 amplitudes than females. In other words, relationships between noise exposure history and AN function could potentially represent sex differences in both variables. In the future, researchers should control for sex differences in noise history exposure and AN function when studying the relationship between noise exposure history and AN function.

Methods for assessing noise exposure must also be considered. In animals, noise exposure can be controlled with a high degree of precision. In humans, however, researchers have largely used self-report measures of noise exposure history, relying on participants’ recollection and subjective assessment of their own noise exposure. While some have employed more rigorous self-report methods, such as detailed structured interviews, to try to better capture the type and amount of noise participants are exposed to, there is little evidence to suggest that these more rigorous approaches result in measures of noise exposure that are more accurate or more sensitive to hearing outcomes or AN function (e.g., Couth et al., 2020; De Poortere et al., 2023; Dewey et al., 2020; Pinsonnault-Skvarenina et al., 2022b). Using more objective measures of prolonged noise exposure – evaluated using a combination of questionnaires and body-worn dosimeters to capture the noise levels and exposure times of participants working in factories – has also failed to find a relationship between noise exposure and AN response amplitudes (Pinsonnault-Skvarenina et al., 2022a; Skoe & Tufts, 2018), though one study did find that noise exposure related to longer ABR WI latencies (Skoe & Tufts, 2018).

A final important factor contributing to the inconsistent relationship between noise exposure history and AN response amplitudes observed between studies is the small effect size calculated across studies (**Figure 4**). Typically, small effect sizes of this kind suggest that larger samples are required to increase statistical power and overcome the probability of a type II error (false negative). We performed a simple a priori power analysis in G*Power to determine the sample size required for the mean noise exposure effect size of r=-0.152 to reach 85% power for a two-tailed test at α=0.05 (Faul et al., 2009; Faul et al., 2007). The results of the power analysis suggested that a sample of 385 participants is needed to reliably capture the effect of noise exposure on AN function. (For comparison, a power analysis for the mean age effect size of r=-0.407 determined that a sample of 51 participants is needed to reach 85% power for a two-tailed test at α=0.05.) Of the studies testing the relationship between noise exposure history and AN function, no single study has a large enough sample (enough statistical power) to confidently assess the relationship. In fact, the studies with the largest samples, including our current CAP study, do not find a relationship between noise exposure and AN function (Couth et al., 2020; Kamerer et al., 2019; Prendergast et al., 2017a; Smith et al., 2019).

Taken together, the evidence for noise-induced effects on AN function in humans is weak, subject to extraneous variables that covary with noise exposure history and AN function and the unknown reliability of self-report assessments of prior noise exposure. The mean effect calculated across studies suggests that any effect of noise exposure history on AN response amplitudes is small, requiring large samples to confidently test and capture the relationship. We can conclude from the results of our own study and our meta-analysis that researchers investigating noise-induced AN damage in humans should be cautious when designing studies, analyzing data, and interpreting results.

### Characteristics of Age-Related and Noise-Induced Auditory Nerve Pathology

The reduced amplitude of the ABR WI following noise exposure has been suggested as a proxy for cochlear synaptopathy based on experiments in noise-exposed animals. These animal studies involved carefully titrated noise and often included histologic confirmation of synapse loss. The presence of such synapse loss in humans cannot be confirmed *in vivo*, and so ABR WI/CAP N1 response amplitudes have been used by many researchers as a proxy for both age-related and noise-induced cochlear synaptopathy. Age-related and noise-induced changes in ABR WI/CAP N1 amplitudes may, however, reflect changes in multiple characteristics of the peripheral auditory system, including cochlear damage, synapse loss, axon degeneration, and even myelin degradation, which is consistent with evidence from human temporal bones from older donors (Makary et al., 2011; Viana et al., 2015; Xing et al., 2012). To date, the specific pathology underlying reduced AN response amplitudes can only be identified with post-mortem histological assessment.

Longitudinal studies are warranted to assess changes in AN function across the lifespan and how such changes in AN function relate to AN structure evaluated post-mortem (for reviews and similar recommendations, see Bramhall et al., 2019; Le Prell, 2019; Shehabi et al., 2022).

### Alternative Methods

The focus of the current systematic review, study, and meta-analysis was on AN suprathreshold response amplitudes. The use of the ABR WI/CAP N1 remains the only *in vivo* method for measuring AN function. While studies to-date have primarily used AN response amplitudes as the sole metric for AN function, other metrics derived from the AN response should be considered for future studies (Harris et al., 2017; McClaskey et al., 2022). Alternative methods besides the AN response have been hypothesized to relate to AN dysfunction, most of which have focused on the loss of synapses. These techniques include behavioral assessments and middle ear reflex measures. In addition, speech perception outcomes, including speech-in-noise and time-compressed speech recognition, have been found to relate to stimulus-evoked AN responses (Harris et al., 2021). The inclusion of these methods is beyond the scope of the current report, but the advantages and disadvantages of these techniques have been reviewed in several papers (Bharadwaj et al., 2019; Bramhall et al., 2019; Harris & Bao, 2022; Le Prell, 2019).

## Conclusion

The systematic review, study, and meta-analyses reported here suggest that age is a robust predictor of AN function, whereas noise exposure history is much less so. Our own study of how AN function relates to age and noise exposure history is consistent with our meta-analyses, finding that CAP N1 response amplitudes decline with age, but are unrelated to noise exposure history. Importantly, our CAP study allowed for the first time a test of how age may interact with noise exposure history to exacerbate age-related deficits in AN function, but we did not find a history of noise exposure to significantly contribute to the age-related deficits in AN function observed in older adults. The evidence from our CAP study and from our meta-analysis suggests that if noise exposure relates to human AN activity, the relationship is small and potentially subject to methodological choices and uncontrolled variables that may covary with AN function and noise-exposure history, including participant age and sex. These results are of clinical importance as AN dysfunction may negatively impact auditory perception and contribute to speech recognition deficits, particularly in the presence of background noise (Bharadwaj et al., 2019; Bramhall et al., 2019; Bramhall et al., 2015; Bramhall et al., 2018a; Grant et al., 2020; Harris et al., 2021).

