## Supplementary material for "Effects of Age and Noise Exposure History on Auditory Nerve Response Amplitudes: A Systematic Review, Study, and Meta-Analysis": MUSC Noise History Questionnaire (from another version of this preprint: version 1, 2024-03-25)

MEDICAL UNIVERSITY OF SOUTH CAROLINA
DEPARTMENT OF OTOLARYNGOLOGY - HEAD & NECK SURGERY
HEARING RESEARCH PROGRAM

### NOISE HISTORY QUESTIONNAIRE

Name________________________________________ Today's Date_______________

Date of Birth________________________________

If more space is needed to answer any of the questions, please use the back of this form.

1. Have you ever worked, or do you now work, in a noisy environment?

   Yes No (Circle one)

   If **NO**, go on to Item #2. If **YES**, describe the environment(s), including military history:

   ____________________________________________________________________

   ____________________________________________________________________

   ____________________________________________________________________

   How many hours a day?______________________________________________

   How many years?___________________________________________________

   How many years, if any, did you wear hearing protection?_______________________

   What type (muffs, cotton, foam plugs)?_____________________________________

2. Have you ever used or do you now use guns? Yes No (Circle one)

   If **NO**, go on to Item #3. If **YES**, describe what types of guns and how often you use(d) them:

   ____________________________________________________________________

   ____________________________________________________________________

   ____________________________________________________________________

   How many years did you use or have you used guns?__________________________

   Did/do you use hearing protection when using guns?

   Never Sometimes Always (Circle one)

   If so, what type?___________________________________________________

3. Do you listen to loud music? Yes No (Circle one)

   If **NO**, go on to Item #4. If **YES**, use the following scale to answer the next two questions:

   soft soft-medium medium medium-loud loud

   How loud is the music when listening through speakers?_______________

   How loud is the music when listening through headphones?____________

4. Have you ever used or do you now use power tools?

Yes No (Circle one)

If **NO**, go on to Item #5. If **YES**, describe what type of power tools you use(d) :

____________________________________________________________

____________________________________________________________

How often did/do you use power tools?________________________________

Did/do you use hearing protection when using power tools?

Never Sometimes Always (Circle one)

If so, what type?___________________________________________

5. Have you ever worked, or do you now work, with farm machinery?

Yes No (Circle one)

If **NO**, go on to Item #6. If **YES**, what type of farm machinery did/do you work with?

____________________________________________________________

____________________________________________________________

How many years have you worked with farm machinery?____________________

Did/do you use hearing protection when using farm machinery?

Never Sometimes Always (Circle one)

If so, what type? ________________________________________________

6. Have you ever been exposed to any sudden loud sound that caused your ears to ring or hurt?

Yes No (Circle one)

If **NO**, go on to Item #7. If **YES**, please describe:____________________

____________________________________________________________

____________________________________________________________

7. Please list any other noise exposure history not listed above and describe in as much detail as possible:________________________________________________

____________________________________________________________

____________________________________________________________

____________________________________________________________
